# Improving annotation propagation on molecular networks through random walks: Introducing ChemWalker

**DOI:** 10.1101/2022.11.04.514505

**Authors:** Tiago C. Borelli, Gabriel Santos Arini, Luís G. P. Feitosa, Pieter C. Dorrestein, Norberto P. Lopes, Ricardo R. da Silva

**Affiliations:** NPPNS, Department of Molecular Biosciences, School of Pharmaceutical Sciences of Ribeirão Preto, University of São Paulo; Collaborative Mass Spectrometry Innovation Center, Skaggs School of Pharmacy and Pharmaceutical Sciences, University of California

## Abstract

Annotation of the mass signals is still the biggest bottleneck for the untargeted mass spectrometry analysis of complex mixtures. Molecular networks are being increasingly adopted by the mass spectrometry community as a tool to annotate large scale experiments. We have previously shown that the process of propagating annotations from spectral library matches on molecular networks can be automated using Network Annotation Propagation (NAP). One of the limitations of NAP is that the information for the spectral matches is only propagated locally, to the first neighbor of a spectral match. Here we show that annotation propagation can be expanded to nodes not directly connected to spectral matches using random walks on graphs, introducing the ChemWalker python library. Similarly to NAP, ChemWalker relies on combinatorial in silico fragmentation results, performed by MetFrag, searching biologically relevant databases. Departing from the combination of a spectral network and the structural similarity among candidate structures, we have used MetFusion Scoring function to create a weight function, producing a weighted graph. This graph was subsequently used by the random walk to calculate the probability of ‘walking’ through a set of candidates, departing from seed nodes (represented by spectral library matches). This approach allowed the information propagation to nodes not directly connected to the spectral library match. Compared to NAP, ChemWalker has a series of improvements, on running time, scalability and maintainability and is available as a stand alone python package. ChemWalker is freely available at https://github.com/computational-chemical-biology/ChemWalker.

## 1 Model description

The MetFusion [1] model uses spectral similarity to inform *in silico* fragmentation prediction. The approach re-calculates the MetFrag score using the following formulation

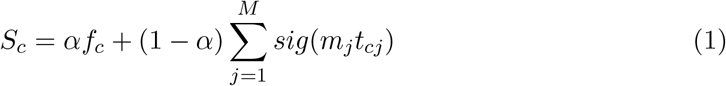

where the index *c* represents each MetFrag candidate, *f*_*c*_ the MetFrag [2] score and the spectral similarity is represented by the spectral library match score *m*_*j*_ for all *j* neighbor nodes, multiplied by the chemical similarity *t*_*cj*_ between MetFrag candidates *c* associated to each neighbor node *j*. The *sig* represents the sigmoid function.

The Network Annotation Propagation (NAP) model uses the MetFusion concept to propagate annotations from spectral library matches through the network topology [3]. The Figure 1 illustrates the propagation concept.

**Figure 1:**
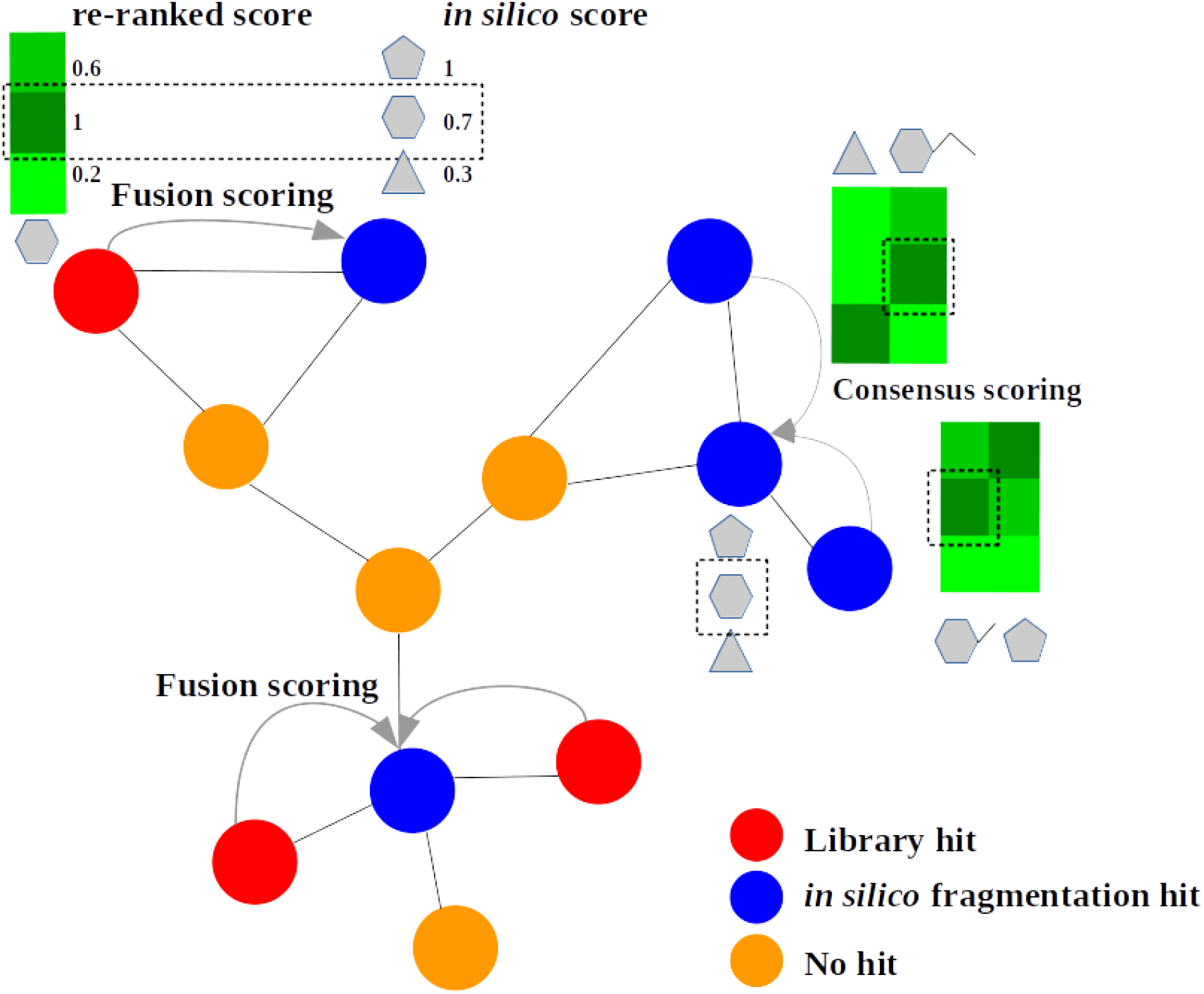
Network Annotation Propagation. The *Fusion* scoring uses the reference from spectral similarity to evaluate structural similarity to candidate structures and re-rank the candidates. The *Consensus* scoring uses the similarity among candidates to retrieve sets of similar candidates.

### 1.1 Structural similarity graph

From NAP’s model one can derive a graph of structural similarity, following the spectral network topology. The Figure 2 illustrates the replacement of the spectra, by its *in silico* fragmentation candidates and the generation of a structural similarity graph, where the edges are weighted by a similarity score, discussed below.

**Figure 2:**
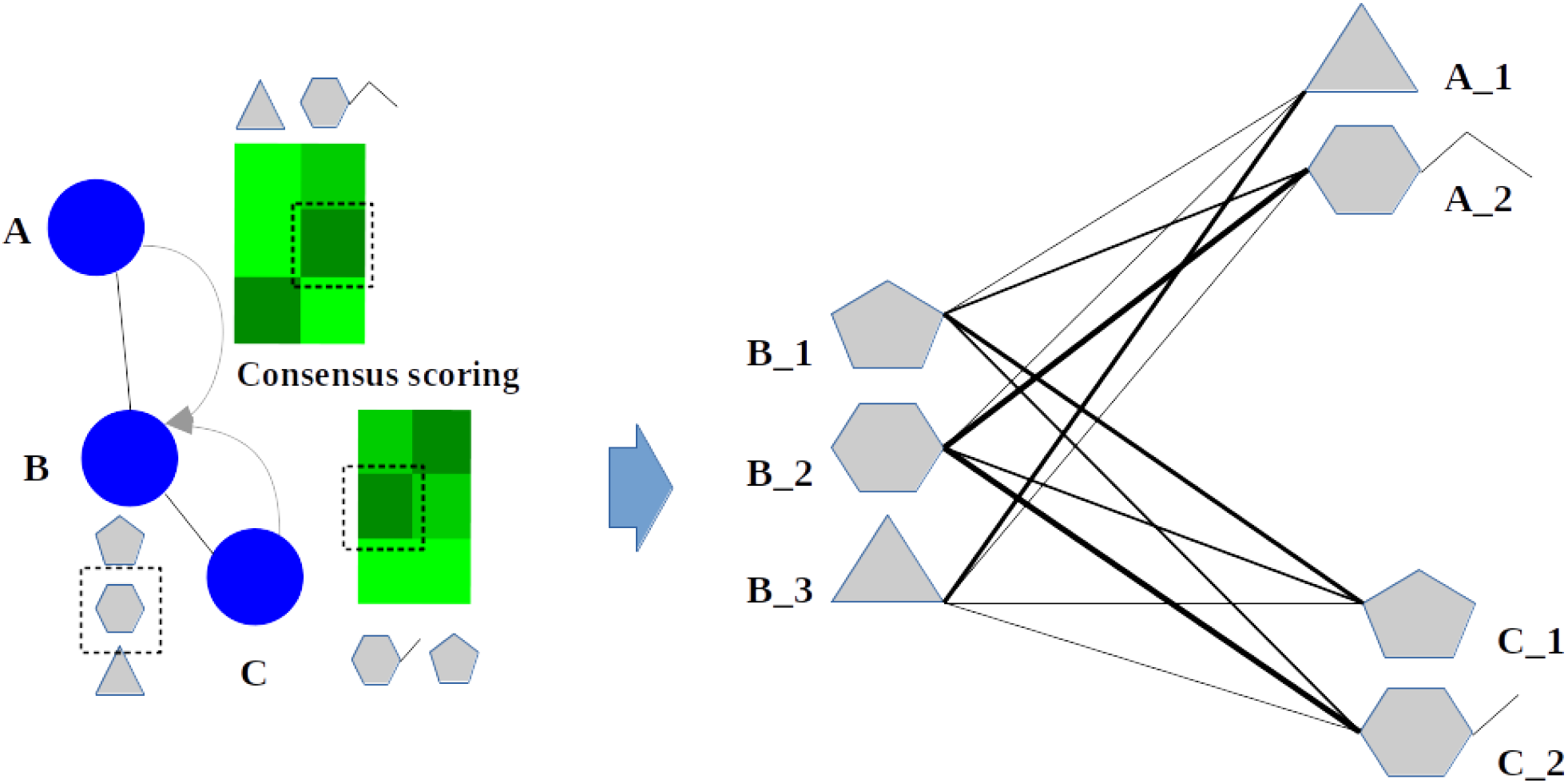
Structural Similarity Graph. The similarity between candidates of the neighbor nodes are used to create a weighted graph.

The *Graph G* = (*N, E*) can be represented by a set of |*N* | = *n* nodes, representing the candidate structures, and |*E*| = *e* edges representing the structural similarity between two given candidates.

### 1.2 Random Walk model

Under the assumption that the spectral similarity observed among the spectra infers the structural similarity of the underlying compounds structures, the Random Walk on graphs selects an initial node *v*_0_, and subsequently an adjacent node *v*_1_ and moves to this neighbor [4]. The walks can be determined by a probabilistic model with of the following form

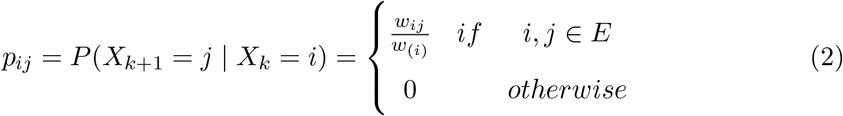

where *w*_(*i*)_ = Σ_*j*∈Γ(*i*)_ *w*_*ij*_, and *i* and *j* represent neighbor nodes in the graph. To maintain the MetFusion score setup, averaging spectral and structural similarities, we applied the following formulation

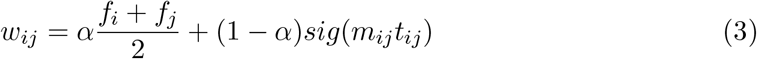

where the indexes *i* and *j* represent two connected nodes, *f*_*index*_ the MetFrag score for each node, the spectral similarity is represented by cosine score *m*_*ij*_ between nodes *i* and *j*, multiplied by the chemical similarity *t*_*ij*_ between MetFrag candidate structures associated to nodes *i* and *j*. The *sig* represents the sigmoid function. The Markov property holds, where conditional on the present, the future is independent of the past, therefore:

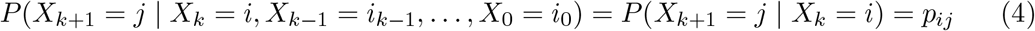

The probability of choosing a given *i-th* candidate structure at a step *k* of the Random Walk is given by the probability vector *p*_*k*_ [5], which at any given step can be obtained by

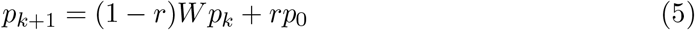

where *W* is the column-normalized adjacency matrix of the graph, *p*_0_ was constructed such that equal probabilities were assigned to the nodes representing ‘seed’ spectral library matches and *r* is the restart probability, which restricts on how far we want the random walker to get away from the start ‘seed’ node(s). The candidate compounds are ranked according to the values in the steady-state probability vector *p*_∞_. This state is obtained at query time by performing the iteration until the change between *p*_*k*_ and *p*_*k*+1_ (measured by the L1 norm) falls below 10^−6^.

### 1.3 Random Walk algorithm

Following the description above the algorithm to compute the candidate probabilities is given by:

#### Algorithm 1

Random Walk on Graphs algorithm

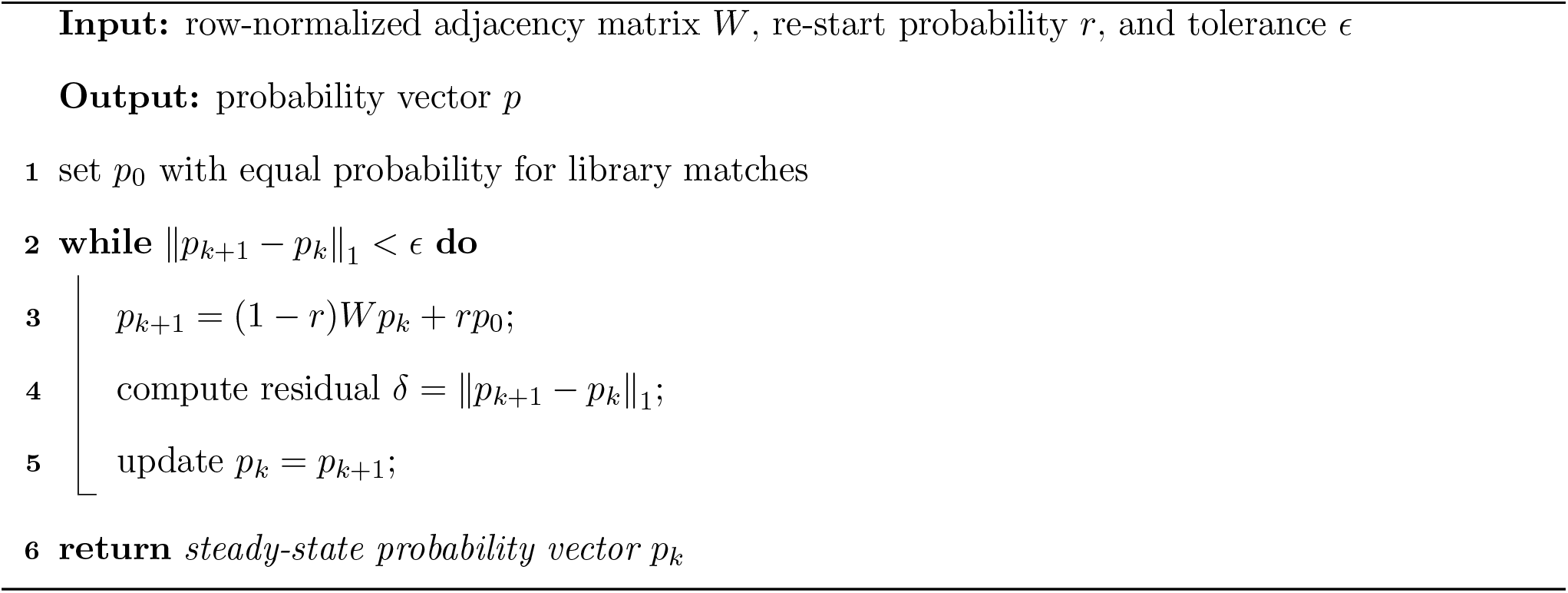

## 2 Benchmarking ChemWalker

In order to benchmark NAP, 5,467 spectra from NIST library were used to compare the propagation scores to MetFrag used on each spectra individually. To extend the benchmark, a subset of 555 spectra, keeping the proportionality of the NAP’s result was selected [3], to compare the previous results with ChemWalker (Figure 3).

**Figure 3:**
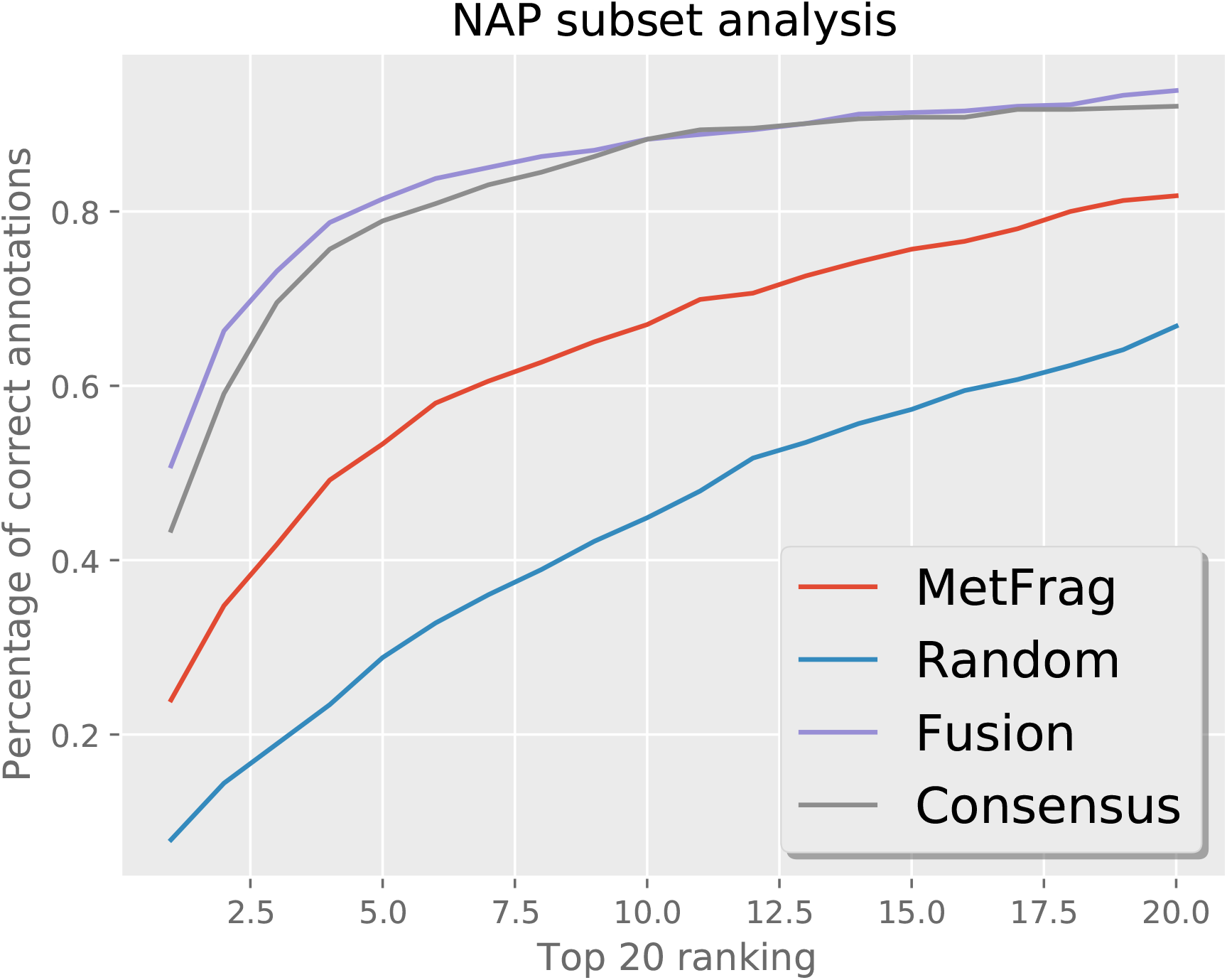
Spectra sub-set from NAP’s validation used to ChemWalker comparison.

First, the molecular fingerprint used for propagation was compared. The RDKit (https://www.rdkit.org/) library was used, and 27 fingerprints were compared. As ChemWalker was designed to work with all spectral library matches as seed nodes, we established 10% of nodes of a given connected component to be the number of seeds in the benchmark dataset (one node for connected components with less than 10 nodes). The fingerprint comparison is shown on Figure 4.

**Figure 4:**
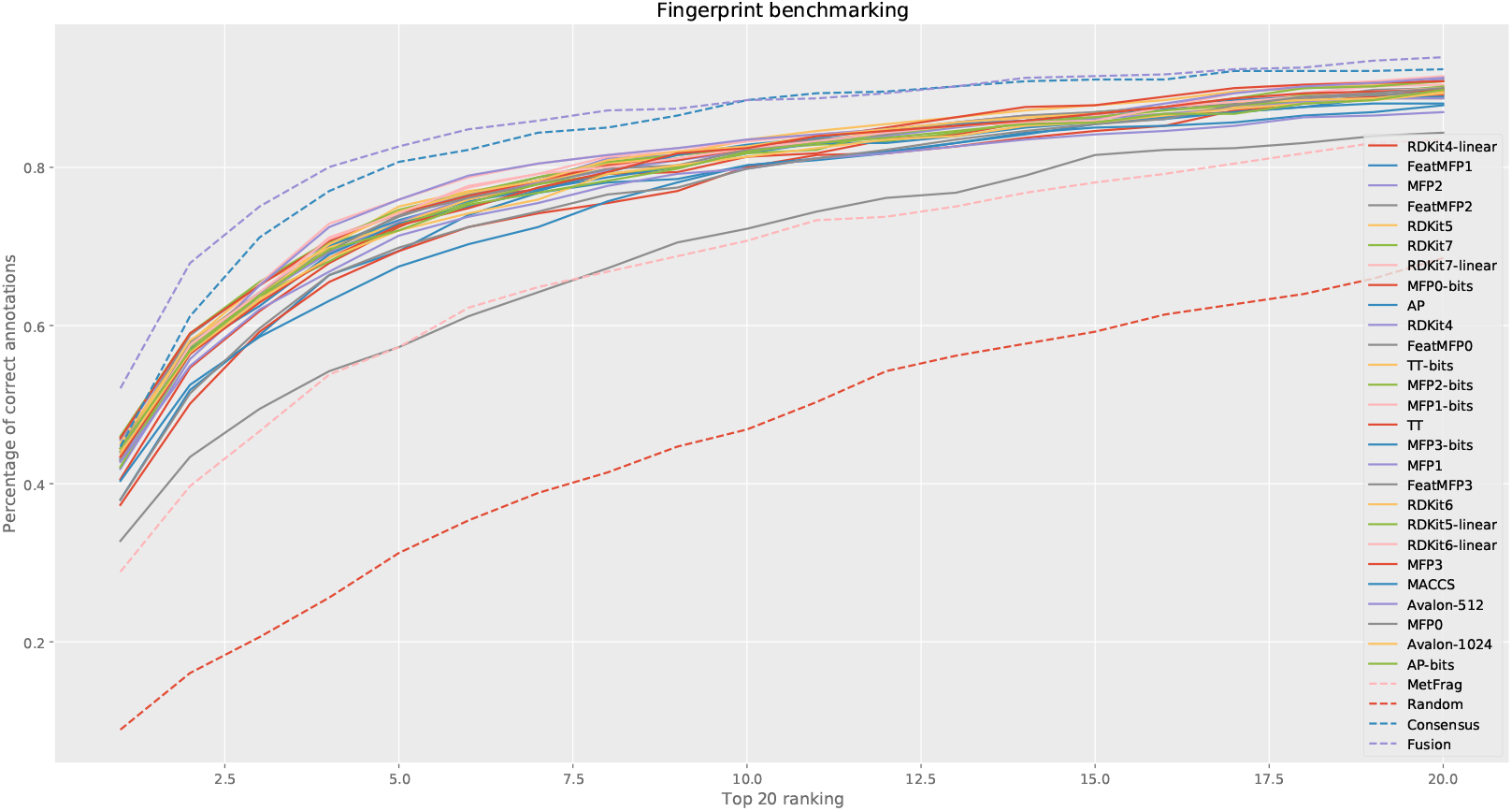
Fingerprint comparison for ChemWalker, using RDKit library.

Despite the improvement over Random and MetFrag, and a higher number of correct candidates ranked in the first three position, comparing *Consensus* scoring with the fingerprints *MFP2, TT-bits, MFP2-bits* and *TT*, the visual inspection shows an overall superiority for *Consensus* and *Fusion* scoring. This was expected, since all neighbors of all nodes act as additional information for these scores, while, for ChemWalker, there are connected components ranging from 346 to 2 nodes, and the only information available comes from seed nodes. Figure 5 shows a connected component of 15 nodes, where 11 were improved over MetFrag. The two *seed* nodes where randomly placed, and we can see an example of improvement even at 3 nodes distance (distance from lower blue border *seed*, node 57 to top green border improved classification, node 159). The benchmark analysis can be found at *benchmark*.*ipynb* on ChemWaker’s git repository.

**Figure 5:**
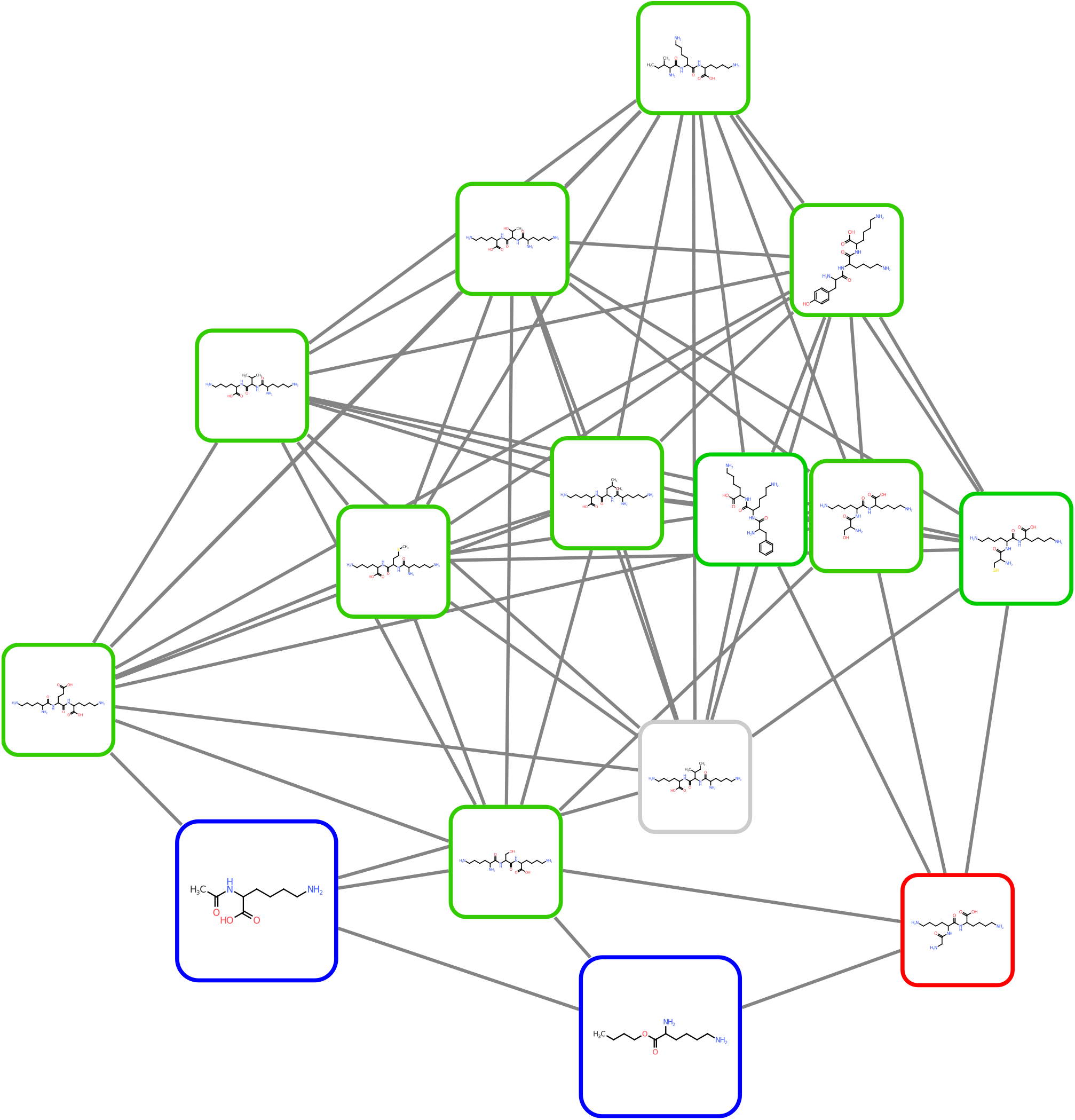
Connected component analysed by ChemWalker. Larger nodes represent *seed* spectral library matches, from where random walks re-start. Green borders represent improvement over MetFrag, red nodes represent loss over MetFrag and grey no ranking alteration.

Carefully inspecting some of the best performing fingerprints, we observe in Figure 6, that the difference between the performance improvement of ChemWalker over MetFrag seems compatible to *Consensus* and *Fusion* scoring.

**Figure 6:**
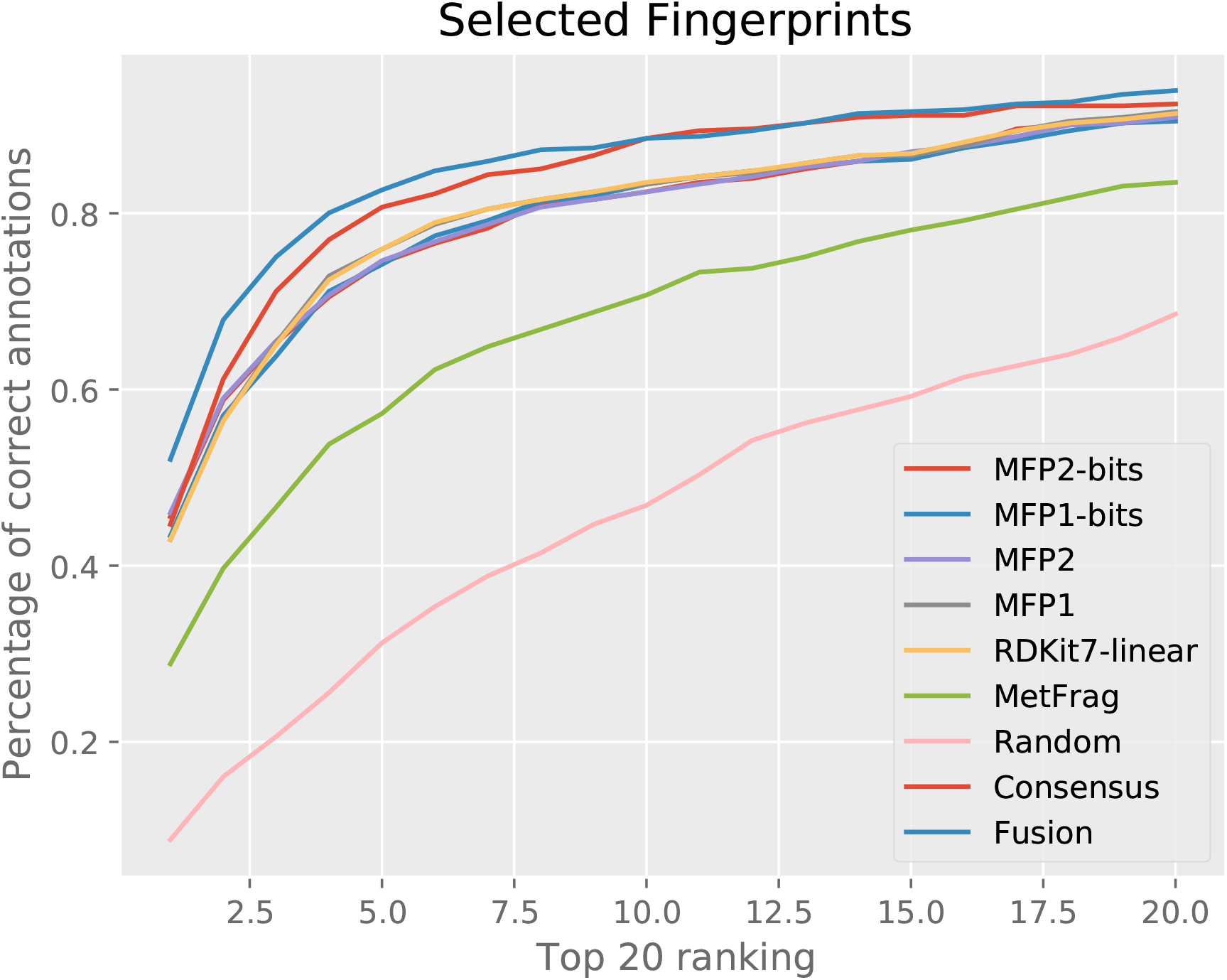
Fingerprint comparison for ChemWalker, using selected RDKit library fingerprints.

Using Chi-square to test the goodness of fit for the number of candidates classified in the 20 first positions, between ChemWalker with different fingerprints and NAP, we can see that some of the best performing fingerprints cannot be statistically differentiated from *Consensus* and *Fusion* scores (Table 1).

**Table 1:**
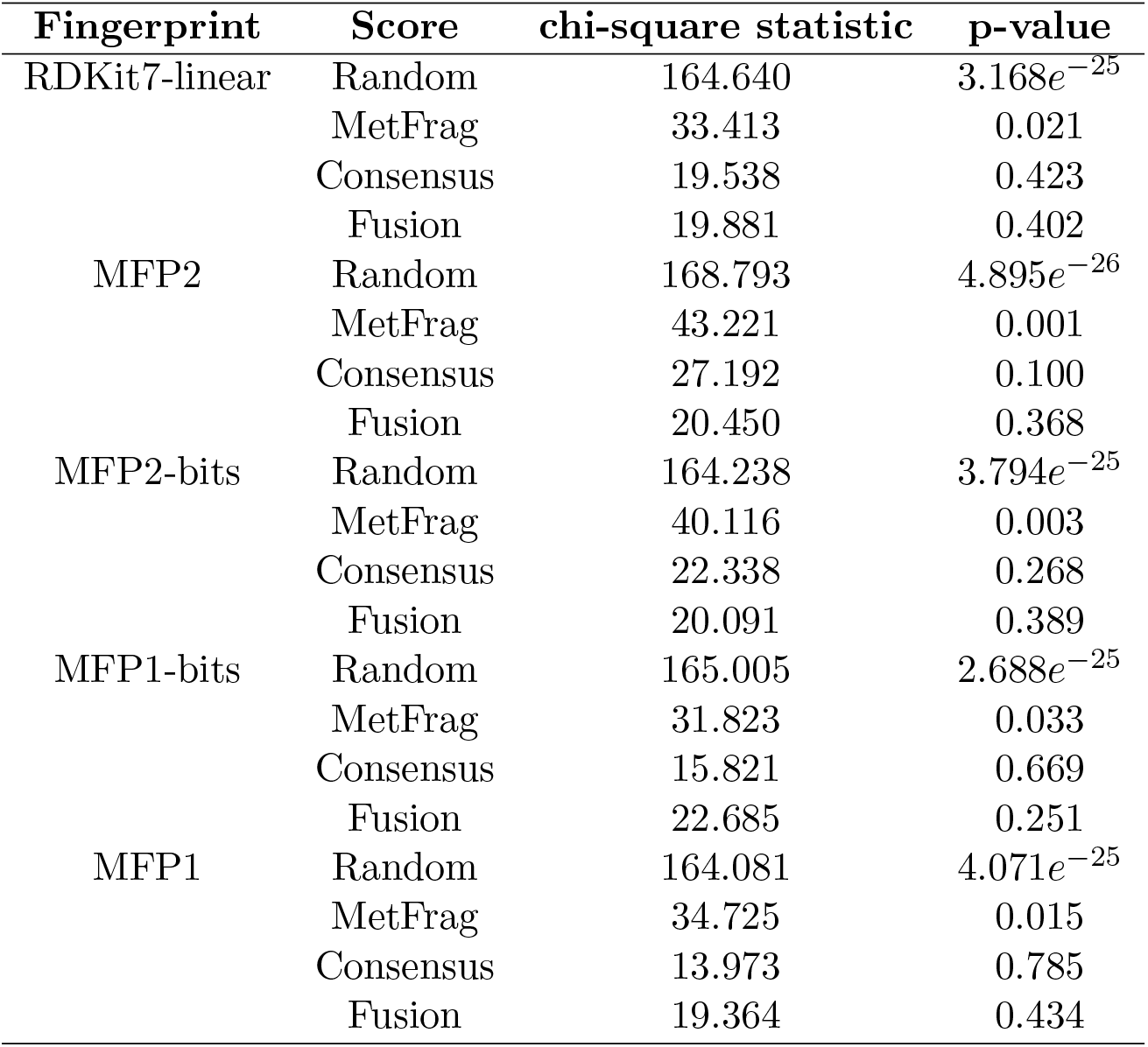
Comparison between ChemWalker, with selected fingerprints, to NAP’s scores.

| **Fingerprint** | **Score** | **chi-square statistic** | **p-value** |
| --- | --- | --- | --- |
| RDKit7-linear | Random | 164.640 | $3.168e^{-25}$ |
|  | MetFrag | 33.413 | 0.021 |
|  | Consensus | 19.538 | 0.423 |
|  | Fusion | 19.881 | 0.402 |
| MFP2 | Random | 168.793 | $4.895e^{-26}$ |
|  | MetFrag | 43.221 | 0.001 |
|  | Consensus | 27.192 | 0.100 |
|  | Fusion | 20.450 | 0.368 |
| MFP2-bits | Random | 164.238 | $3.794e^{-25}$ |
|  | MetFrag | 40.116 | 0.003 |
|  | Consensus | 22.338 | 0.268 |
|  | Fusion | 20.091 | 0.389 |
| MFP1-bits | Random | 165.005 | $2.688e^{-25}$ |
|  | MetFrag | 31.823 | 0.033 |
|  | Consensus | 15.821 | 0.669 |
|  | Fusion | 22.685 | 0.251 |
| MFP1 | Random | 164.081 | $4.071e^{-25}$ |
|  | MetFrag | 34.725 | 0.015 |
|  | Consensus | 13.973 | 0.785 |
|  | Fusion | 19.364 | 0.434 |

